# Natural variation in rice *mitogen-activated protein kinase 4* contributes to increased photosynthetic rate under field conditions

**DOI:** 10.64898/2026.03.06.710232

**Authors:** Tadamasa Ueda, Shunsuke Adachi, Kazuhiko Sugimoto, Miki H. Maeda, Utako Yamanouchi, Ritsuko Mizobuchi, Yojiro Taniguchi, Tadashi Hirasawa, Toshio Yamamoto, Junichi Tanaka

**Affiliations:** Institute of Crop Science, National Agriculture and Food Research Organization (NARO), Tsukuba, Ibaraki 305-8518, Japan; Tokyo University of Agriculture and Technology, Fuchu, Tokyo 183-8509, Japan; National Institute of Animal Health, NARO, Tsukuba, Ibaraki 305-0856, Japan; Okayama University Institute of Plant Science and Resources, Kurashiki, Okayama 710-0046, Japan; NARO Headquarters, Tsukuba, Ibaraki 305-8518, Japan; University of Tsukuba, Graduate School of Science and Technology, Tsukuba, Ibaraki 305-8577, Japan; Tsukuba Operations Unit 1, NARO, Tsukuba, Ibaraki 305-8518, Japan

**Keywords:** Chromosome segment substitution lines, CRISPR/Cas9, map-based cloning, near-isogenic line, *OsMPK4* (Os10g0533600), photosynthetic rate, *qHP10*, rice (*Oryza sativa* L.), stomatal conductance (9 words)

## Abstract

Improving rice (*Oryza sativa* L.) yield requires a balanced enhancement of both sink size and source capacity. While many QTLs for sink size have been identified, only a few are known for source capacity, which is essential for achieving high yield. Here we identified *qHP10* as a major QTL for increased photosynthetic rate by using chromosome segment substitution lines derived from a cross between the high-yielding *indica* cultivar Takanari and the average-yielding *japonica* cultivar Koshihikari. High-resolution mapping combined with CRISPR/Cas9-induced mutagenesis revealed that the causative gene underlying *qHP10* is *Mitogen-Activated Protein Kinase 4* (*OsMPK4*). A near-isogenic line carrying the *OsMPK4^Takanari^* allele (NIL-*OsMPK4*) had a 15–25% higher photosynthetic rate than Koshihikari. NIL-*OsMPK4* also had higher stomatal conductance than Koshihikari but similar stomatal pore size and density, indicating that increased stomatal aperture increases photosynthetic rate. This enhancement is likely attributable to the down-regulation of *OsMPK4* expression, which increases stomatal conductance and thus promotes CO_2_ uptake. Our findings demonstrate that *OsMPK4* is a promising genetic target for increasing source capacity and, potentially, rice yield through molecular breeding. (175 words)

## Introduction

As the global population continues to grow, increasing food production has become a critical challenge. Rice (*Oryza sativa* L.) is one of the most important food crops, feeding nearly half of the world’s population; therefore, genetic improvement to enhance grain yield remains a major challenge in rice breeding. To achieve higher yield, the sink size (the size and number of harvestable organs), source capacity (the capacity for photosynthetic carbon assimilation), and efficiency of carbon partitioning and translocation must all be improved. Although rice breeding has steadily increased yields over the decades, these gains have largely resulted from modifications in carbon partitioning or sink size rather than from improvements in photosynthesis (Orr *et al*., 2017; Ueda *et al*., 2025). Thus, enhancing photosynthetic capacity remains a promising route to further yield gains (Evans, 2013; Sharwood, 2016; Chen *et al*., 2021; Wei *et al*., 2022).

Many attempts have been made to improve leaf photosynthesis through genetic engineering (Evans, 2013; Sharwood, 2016; Kubis and Bar-Even, 2019). For example, overexpression of *HIGHER YIELD RICE* (*HYR*), a transcription factor, increases photosynthetic rate and yield of rice under drought and high-temperature stress conditions (Ambavaram *et al*., 2014). Knockout of *NEGATIVE REGULATOR OF PHOTOSYNTHESIS 1* (*NRP1*) induces the expression of photosynthesis-related genes, enhancing leaf photosynthesis and biomass production in rice (Chen *et al*., 2021). Overexpression of the transcription factor *Dehydration-Responsive-Element-Binding Protein 1C* (*OsDREB1C*) promotes early flowering, enhances photosynthesis, and improves nitrogen use efficiency, resulting in higher yield in both rice and wheat (*Triticum aestivum* L.) (Wei *et al*., 2022). These results indicate that increased photosynthetic rate may lead to an increase in rice yield and/or biomass, but these genetic improvements have not yet been widely adopted in practical rice cultivars.

Using natural genetic resources in rice is also a promising approach to improving its photosynthetic rate (Cook and Evans, 1983; Sasaki and Ishii, 1992; Kanemura *et al*., 2007; Jahn *et al*., 2011). The high-yielding *indica* rice cultivar Takanari (Imbe *et al*., 2004) has one of the highest photosynthetic rates in the flag leaf among a wide range of rice genetic resources (Xu *et al*., 1997; Kanemura *et al*., 2007). Our research group has conducted studies to identify the underlying genetic factors using genetic mapping populations such as backcross inbred lines and chromosome segment substitution lines (CSSLs) derived from a cross between Takanari and average-photosynthesis *japonica* cultivar Koshihikari (Takai *et al*., 2013, 2014). Several quantitative trait loci (QTLs) for photosynthetic rate have been identified, including *Green for Photosynthesis* (*GPS*) and *High Photosynthesis on Chromosome 10* (*qHP10*), in which the Takanari allele improved photosynthesis (Takai *et al*., 2013; Adachi *et al*., 2019). The *GPS* gene was found to be identical to *Narrow Leaf 1* (*NAL1*) (Qi *et al*., 2008) and *SPIKE* (Fujita *et al*., 2013), which encode a serine protease that degrades the TOPLESS-related corepressor (Li *et al*., 2023). Plants carrying the Takanari allele at *GPS* showed an increased number of mesophyll cells and greater leaf thickness, resulting in higher photosynthetic rate (Takai *et al*., 2013). Furthermore, Yamashita *et al*. (2022) reported that a near-isogenic line (NIL) carrying the Takanari allele of *qHP10* had higher flag leaf photosynthesis, stomatal conductance, biomass, and grain production than Koshihikari. However, the causal gene underlying *qHP10* was not identified.

Mitogen-activated protein kinase (MAPK) cascades are conserved signaling pathways that regulate diverse physiological processes; in plants, these include photosynthesis and stomatal movement (Nakagami *et al*., 2005; Rodriguez *et al*., 2010). In *Arabidopsis* and tobacco (*Nicotiana tabacum* L.), *MPK4* is known to be involved in the regulation of stomatal aperture (Hõrak *et al*., 2016; Lin and Chen, 2018), but until now there has been no report linking any monocot homolog of *MPK4*, including *OsMPK4*, to photosynthetic traits.

In this study, we characterized the *qHP10* QTL. High-resolution mapping and phenotypic analysis of mutants induced by clustered regularly interspaced short palindromic repeats (CRISPR)/CRISPR-associated protein 9 (Cas9) revealed that the causal gene underlying *qHP10* is *OsMPK4*. A NIL carrying the *OsMPK4^Takanari^*allele in a Koshihikari genetic background (NIL-*OsMPK4*) had a 15–25% higher photosynthetic rate than Koshihikari, primarily owing to increased stomatal conductance resulting from reduced *OsMPK4* expression and increased stomatal aperture, without negative pleiotropic effects on yield. Protein structure prediction and sequence comparison suggested that the absence of a TATA-binding-protein interaction site in *OsMPK4^Takanari^* may reduce mRNA expression, thereby promoting larger stomatal aperture. These findings highlight *OsMPK4^Takanari^* as a promising genetic resource for future rice breeding programs aimed at enhancing source capacity and improving yield potential.

## Materials and methods

### Plant growth condition

Field experiments were conducted in a paddy field at NARO Kannondai Field, Ibaraki, Japan (36°03′N, 140°11′E) from 2013 to 2024, and at the University Farm of Tokyo University of Agriculture and Technology (hereafter Fuchu Field), Tokyo, Japan (35°41′N, 139°29′E) from 2014 to 2022.

In NARO Kannondai Field, seeds were sown in nursery boxes, and 30-day-old seedlings were transplanted at a density of one seedling per hill, at a spacing of 15 cm between hills and 30 cm between rows. Basal fertilization consisted of slow-release fertilizer at 8 kg N (as urea), 8 kg phosphorus (P_2_O_5_), and 8 kg potassium (K_2_O) per 10 a.

In Fuchu Field, seeds were sown and seedlings were transplanted as above. Basal fertilization consisted of slow-release fertilizer applied at 3 kg N (as urea), 6 kg phosphorus (P_2_O_5_) and 6 kg potassium (K_2_O) per 10 a.

### Plant materials and map-based cloning

CSSL line SL1235 carries a segment of Takanari chromosome 10 (approximately 12.4 Mb) in the genetic background of Koshihikari (Adachi *et al*., 2019). SL1235 was crossed with Koshihikari to produce F_1_ progeny, which were self-pollinated to obtain F_2_ seeds. For initial mapping of *qHP10*, 476 F_2_ seedlings were grown in Fuchu Field. Plants with recombination events near the *qHP10* region were identified using region-specific markers RM25699 and RM25840. F_3_ lines derived from selected F_2_ plants were further classified into two types: those homozygous for a recombinant chromosome segment (R; recombinant) and those homozygous for a nonrecombinant segment (IC; internal control). Those F_3_ lines were transplanted and evaluated for photosynthesis-related traits. As controls, we used Koshihikari and NIL10, which carries a 2.8 Mb Takanari chromosome fragment including *qHP10* in the Koshihikari genetic background (Adachi *et al*., 2019; Yamashita *et al*., 2022).

For fine mapping, 2800 F_2_ seedlings were grown in NARO Kannondai Field. Plants with recombination events near the *qHP10* region were selected by using the region-specific DNA markers RM25771 and RM25775. We also grew F_3_ lines obtained from the selected F_2_ plants and then selected two types of F_3_ plants: one type homozygous for a recombinant chromosome segment (R; recombinant) and the other homozygous for a nonrecombinant (IC) segment. Selected F_3_ lines were transplanted into Fuchu Field for evaluation of photosynthesis-related traits.

We also developed NIL-*OsMPK4,* which carries a <80.2-kb Takanari chromosome segment containing *qHP10* in the genetic background of Koshihikari, and analyzed its photosynthetic and agronomic properties in NARO Kannondai Field.

### DNA marker analysis

Total DNA was extracted according to Monna *et al*. (2002). All DNA markers in this work were PCR-based markers (Supplementary Table S1). PCR amplification used 35 cycles of 95℃ for 30 s, 50–60℃ for 1 min, and 72℃ for 1 min. Cleaved amplified polymorphic sequence (CAPS) and derived CAPS (dCAPS) markers were digested with the respective restriction enzyme (Supplementary Table S1), and all PCR products were separated by electrophoresis in 3% agarose gel and visualized with ethidium bromide staining.

### Gas exchange measurements for photosynthesis

Leaf photosynthetic rate and stomatal conductance were measured in NARO Kannondai Field and Fuchu Field. using a portable gas exchange system (LI-6400XT; LI-COR Inc., Lincoln, NE, USA). Measurements were conducted on flag leaves at the full heading stage (3–7 days after flowering) between 09:00 and 12:00 on clear days. The chamber conditions were set to saturated light of 2,000 μmol photon m^−2^ s^−1^ provided by red/blue light-emitting diodes, a CO_2_ concentration of 400 μmol mol^−1^, a mean leaf-to-air vapor pressure difference of 1.4 kPa, and a leaf temperature of 30°C.

We also measured the photosynthetic rate under varying light intensities (0–2,000 μmol m^−2^ s^−1^) and CO_2_ concentrations (0-1,000 μmol mol^−1^) in the paddy field. In these measurements, the chamber conditions were set to 2,000 μmol photon m^−2^ s^−1^ of light, a CO_2_ concentration of 400 μmol mol^−1^, a mean leaf-to-air vapor pressure difference of 1.4 kPa, and a leaf temperature of 30 °C. The maximal rate of ribulose-1,5-bisphosphate (RuBP) carboxylation by Rubisco (*V*_cmax_) and the maximal electron transport rate (*J*_max_) were estimated by fitting the Farquhar–von Caemmerer–Berry (FvCB) model (Farquhar *et al*., 1980) using a model-fitting utility implemented in Microsoft Excel ver. 2.95 (Gregory *et al*., 2021).

SPAD value, an index of relative chlorophyll content, was measured using a SPAD-502 chlorophyll meter (Konica Minolta Japan, Inc., Tokyo, Japan).

### Development of CRISPR/CAS9-mediated mutant plants

We followed a previous report (Mikami *et al*., 2016) to edit *OsMPK4* by using CRISPR/CAS9 technology. Two targets were designed in the ORF of *OsMPK4* by using a web tool, CHOPCHOP (https://chopchop.cbu.uib.no): 5′**-**TCAAGGGGATGGGGACGCACG**-**3′ (target1) and 5′-ACAACCATATCGATGCCAAGC-3′ (target2). These oligonucleotides, each containing a protospacer adjacent motif (PAM), were annealed with the antisense sequence. The annealed oligonucleotides were ligated between the *Asc*I and *Eco*RV sites of the pZH_OsU6gRNA_MMCas9 binary vector (Mikami *et al*., 2016).

*Agrobacterium*-mediated transformation of rice was conducted as described previously (Toki, 1997). The seedlings were transplanted into 4-L pots filled with Bonsol No.2 (Sumitomo Chemical Co, Ltd., Osaka, Japan), and the room temperature was 28℃. To screen the T_0_ mutant plants, we conducted direct sequencing of PCR fragments containing target1 or target2. The nucleotide sequences of the amplification primers are listed in Supplementary Table S1. All the mutant plants were grown in an isolated greenhouse to comply with biosafety regulations for genetically modified rice. Screened mutant plants were evaluated for photosynthesis-related traits at the full heading stage.

### Prediction and analysis of 3D-structures of gene products

Three-dimensional structures were predicted by AlphaFold 2.3.0 (DeepMind; https://github.com/deepmind/alphafold), and the top-ranked structure (designated “ranked_0”) according to the AlphaFold score (pLDDT) was selected for each sequence. Output structures were visualized and analyzed by MOE 2024.06 (Chemical Computing Group, Montreal, QC, Canada; https://www.chemcomp.com).

### Observation of stomatal anatomy

For observation of stomatal density and pore length, the middle part of flag leaves was fixed in solution containing (v/v) 5% formalin, 5% acetic acid, and 63% ethyl alcohol in distilled water. The abaxial surfaces of the fixed leaves were photographed under a scanning electron microscope (VE-8800; Keyence, Osaka, Japan). Stomatal number per unit area was counted using a computer installed with original software (Adachi *et al*., 2017). The length of stomatal pores was analyzed with ImageJ software (National Institutes of Health, Bethesda, MD, USA).

### RNA isolation and expression analysis of *OsMPK4*

Total RNA was extracted from flag leaf tissues using an RNeasy Mini Kit (Qiagen, Venlo, the Netherlands). First-strand cDNA was synthesized by using an iScript gDNA Clear cDNA Synthesis Kit (Bio-Rad, Hercules, CA, USA). Quantitative real-time PCR (qRT-PCR) was performed by using Thunderbird SYBR qPCR Mix (Toyobo, Osaka, Japan) and *OsMPK4*-specific primers (Supplementary Table S1) on a ViiA7 system (Applied Biosystems, Waltham, MA, USA). Five biological replicates were used, with four technical repeats for each biological replicate. The *UBQ2* gene (Os02g0161900) from rice was used as an internal standard (Supplementary Table 1).

### Agronomic trait evaluation

Field trials were conducted at NARO Kannondai Field. Heading date was defined as the day when the first panicle emerged. Plant height, panicle length, and panicle number were measured 14–21 days after heading. Plants were harvested 40 days after heading and air-dried for 2 weeks. Paddy rice (i.e., unhusked rice) was hulled and referred to as “ungraded brown rice,” and the ungraded brown rice that passed through a 1.8 mm sieve was classified as “graded brown rice”. The following traits were also determined: grain number per panicle, fertility ratio (fertile grains per panicle/total grain number per panicle), total plant weight, harvested paddy rice weight, weights of ungraded and graded brown rice, and 1000-grain weight of graded brown rice. “Perfect grains” were defined as grains excluding damaged, dead, immature, or foreign grains and foreign substances. The perfect grain ratio (number of perfect grains/number of graded brown rice grains) was quantified using a grain discriminator (RGQl100B; Satake Inc., Higashi Hiroshima, Japan).

For the assessment of resistance to bacterial blight caused by *Xanthomonas oryzae* pv. *oryzae*, flag leaves of 14-week-old plants in NARO Kannondai Field were inoculated as previously described (Mizobuchi *et al*., 2002). A bacterial suspension (OD_600_=0.25) of *X. oryzae* pv. *oryzae* race 1 (T7174, MAFF311018) was used for the inoculation, and lesion length was scored 20 days after inoculation.

## Results

### Map-based cloning and genetic validation of *qHP10*

SL1235, which is homozygous for the *qHP10^Takanari^* allele, had significantly higher leaf photosynthetic rate and stomatal conductance than Koshihikari (homozygous for the *qHP10^Koshihikari^*allele), while internal CO_2_ concentration was similar between the two lines (Supplementary Fig. S1). The plants heterozygous at the *qHP10* locus displayed intermediate phenotypes (Supplementary Fig. S1). From the phenotypic evaluation of 476 plants derived by self-pollination of heterozygous NIL-*qHP10*, we determined that *qHP10* is located between markers RM25771 and RM25775 (Fig. 1A). In a segregating population of 2,800 plants derived from a cross between SL1235 and Koshihikari, we selected 18 F_1_-derived F_2_ recombinant lines between RM25771 and RM25775 for further analysis (Fig. 1B). Photosynthetic analysis of homozygous plants in the F_2_ generation delimited the candidate region of *qHP10* to a 14.5-kb region between DNA markers E and I (Fig. 1B, Supplementary Fig. S2). According to the Rice Annotation Project Database (RAP-DB; https://rapdb.dna.affrc.go.jp) (Sakai *et al*., 2013), this region contains three genes: Os10g0533600, Os10g0533800, and Os10g0533900 (Fig. 1B). Os10g0533800 encodes concanavalin A-like lectin/glucanase, subgroup domain containing protein, and this gene did not express in flag leaf (RAP-DB). Os10g0533900 encodes a protein of unknown function (RAP-DB). Os10g0533600 encodes OsMPK4 (Fig. 1B), and previous studies in *Arabidopsis* and tobacco have shown that *MPK4* is involved in regulating stomatal aperture (Hõrak *et al*., 2016; Lin and Chen, 2018). Therefore, we considered *OsMPK4* to be the most likely candidate gene, which is further supported by the fact that it is expressed in flag leaf (RAP-DB).

**Fig. 1.**
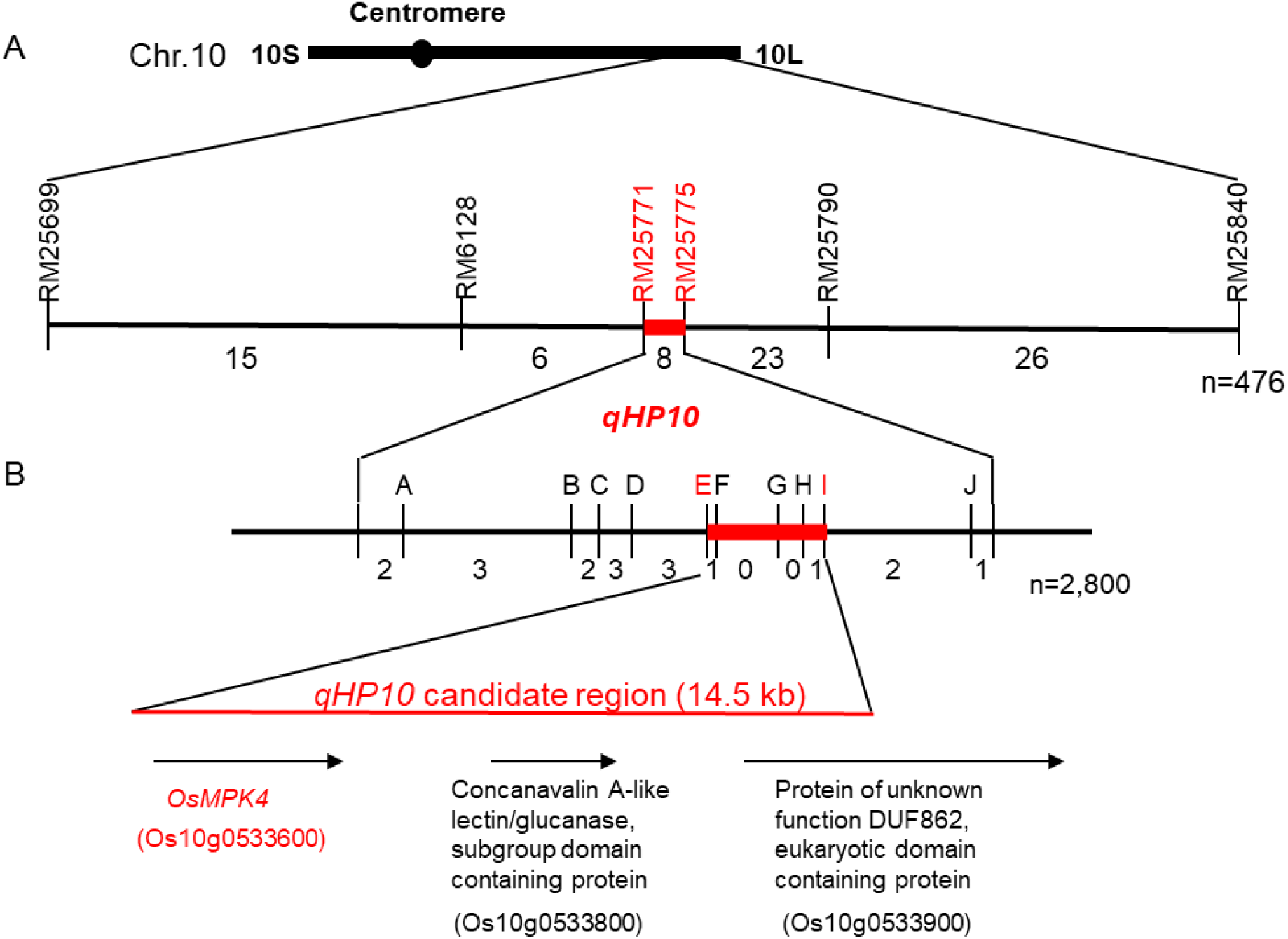
Map-based cloning of *qHP10.* (A) Small-scale mapping of *qHP10*. The names of the DNA markers used for mapping are shown above the horizontal line, and the numbers of recombinants between molecular markers are shown below the line. (B) Fine mapping of *qHP10*. Arrows indicate the genes in the candidate region.

We developed *OsMPK4^Koshihikari^* knockout plants by using CRISPR/CAS9 genome editing. Among 92 T_0_ plants generated, 37 carried mutations around target1 in exon 1, while no mutations were induced near target2 in exon 2 (Fig. 2A). Among these T_0_ plants, lines #37 and #45 were heterozygous, carrying both a wild-type allele and a frame-shift mutation (1-bp deletion) in the *OsMPK4^Koshihikari^* allele. Line #18 was heterozygous for the wild-type allele and a 3-bp deletion, whereas line #50 was heterozygous for 15-bp and 39-bp deletions (Fig. 2A). In the T_1_ generation, we were unable to obtain plants homozygous for the 1-bp deletion from lines #37 and #45, likely owing to lethality. In contrast, from the T_1_ progeny of lines #18 and #50, we obtained plants homozygous for 3-bp, 15-bp, and 39-bp deletions (hereafter referred to as 3bpDel, 15bpDel, and 39bpDel, respectively). Photosynthetic rate in the five genome-edited lines was significantly higher than in control plants (Fig. 2B). Stomatal conductance in the mutants was generally higher than in the control, with only one mutant (3bpDel) showing a statistically significant increase compared with the control (Fig. 2C).

**Fig. 2.**
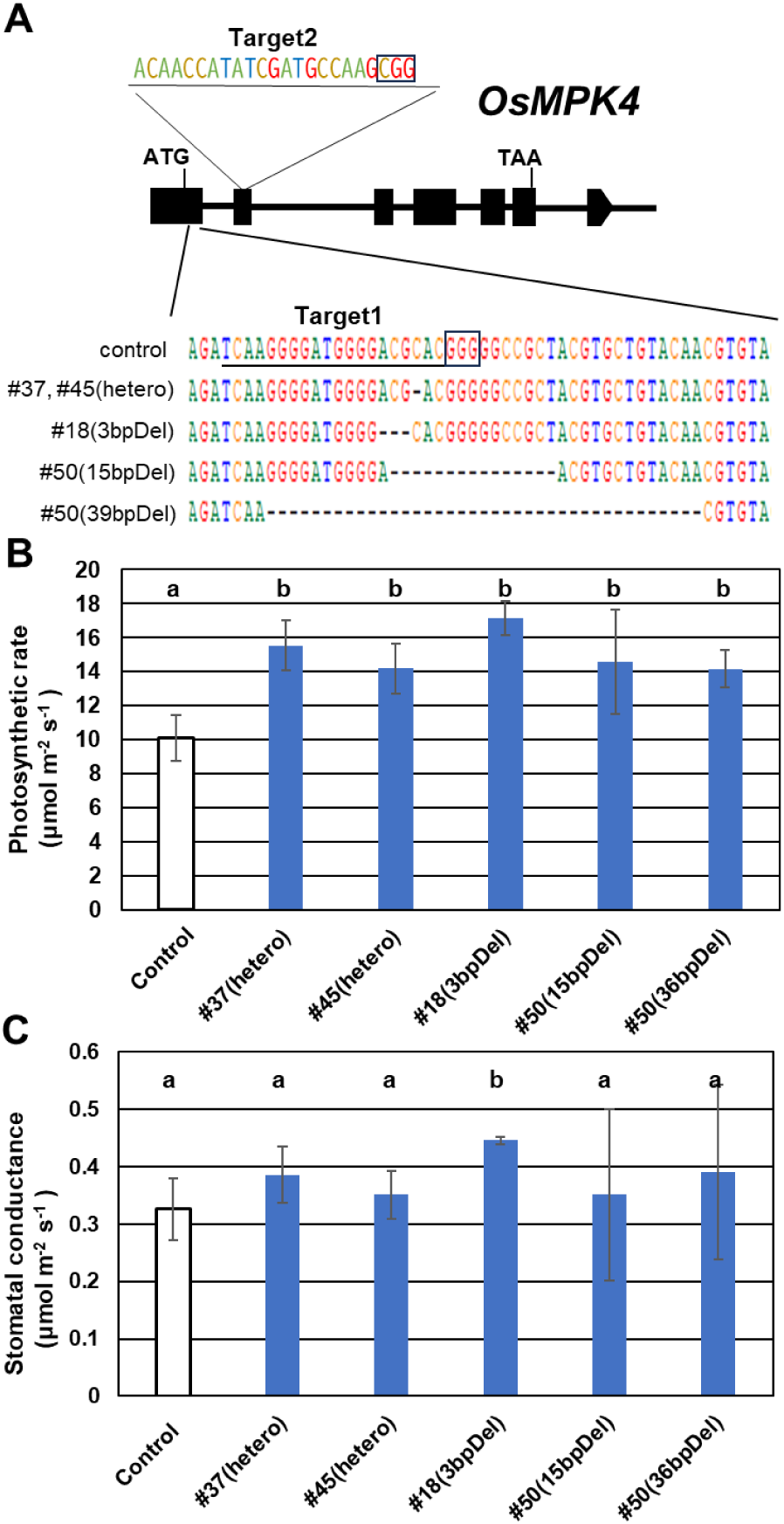
Characterization of *OsMPK4* mutants produced by genome editing. (A) Location of target1 and target2 sequences (underlined) and sequencing result of mutants obtained using CRISPR/CAS9 technology. Each protospacer-adjacent motif (PAM) is marked with a square frame. All of the recovered mutants had mutations around the target1 sequence. (B, C) Comparison of the photosynthetic rate (B) and stomatal conductance (C). Error bars indicate SD (*n*=6). Values marked with the same letter are not significantly different at *P*<0.05 by LSD test.

To elucidate the effects of 3bpDel, 15bpDel, and 36bpDel, we used AlphaFold 2.3.0 to predict four protein structures (representing the three mutants and the wild type). Twenty-five structures provided for each sequence were evaluated by the root mean square deviation (RMSD) for all structures in each set (0.370 Å, 0.572 Å, 0.852 Å, and 1.009 Å for wild type, 3bpDel, 15bpDel, and 39bpDel, respectively). Because all models for each sequence were very similar, the top-ranked structure (designated “ranked_0”) for each one was used for further analysis. The residues lacking from the deletion mutants were all within the first 20 N-terminal residues. The protein parts from Asn31 to the C terminus, which were the same in all four sequences, were predicted to have the same folding (Fig. 3A). The six residues until Val32 were predicted to form a similar β-sheet structure despite their different sequences; furthermore, the wild type and 3bpDel were predicted to have another β-sheet structure prior to this region (Fig. 3B). Comparing the structures of the wild type and 3bpDel, missing Thr21 in the sequences could be location of some residues, especially for Glu15, Lys17, and Met19. From the view in the direction of the green arrow in Fig. 4A, the side-chains of the three residues were on opposite sides of the main chains in the wild type vs. 3bpDel (Fig. 4B). The differences in side-chain positions could lead to a large change in the molecular surface of the site (Fig. 4C, D).

**Fig. 3.**
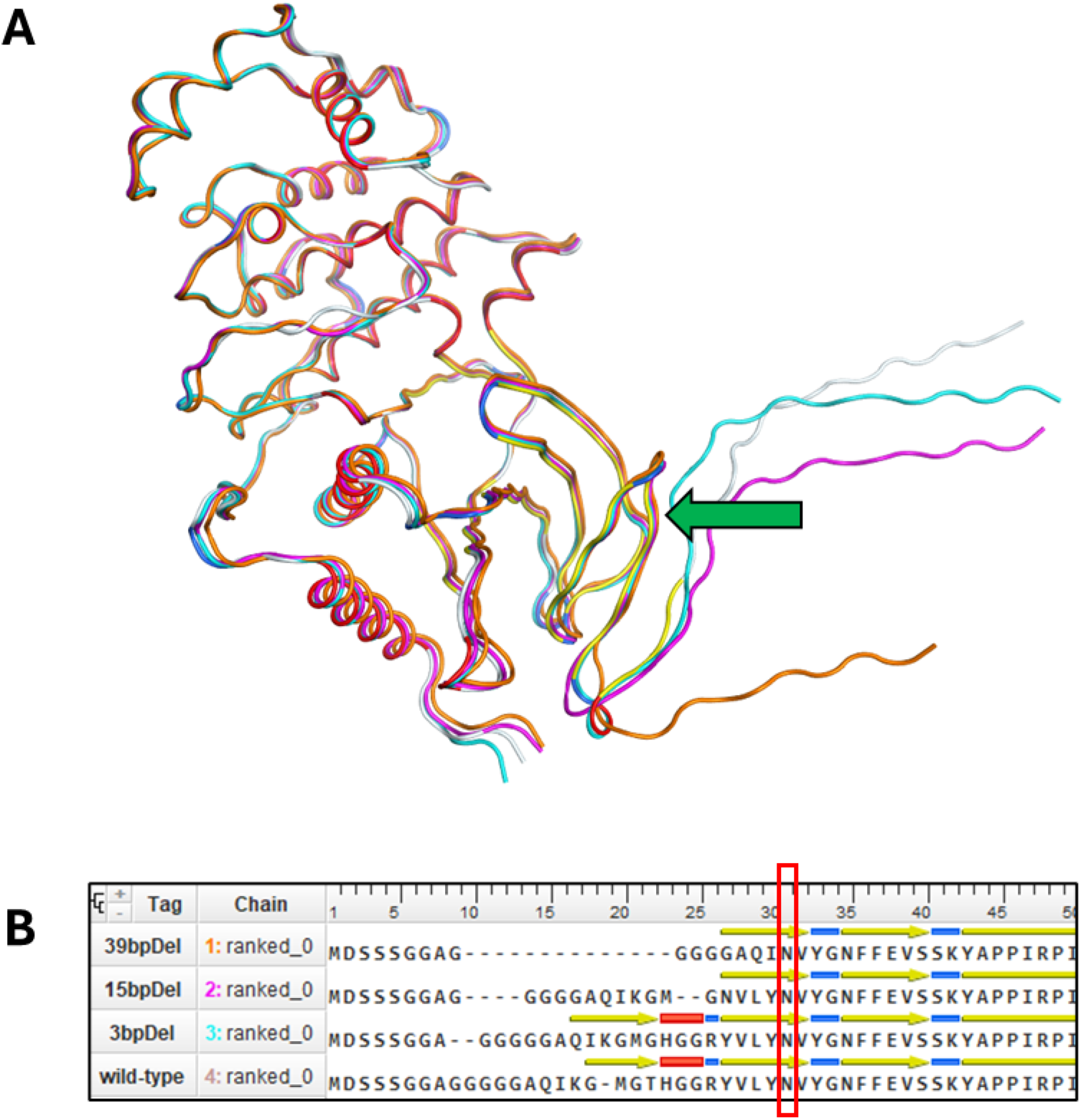
Comparison of four modeled structures of MAPK4: wild type, 3bpDel, 15bpDel, and 39bpDel. (A) Overall superposition of modeled structures of wild type (color-coded according to the secondary structure; i.e., red, yellow, and blue are α-helices, β-sheets, and turns, respectively), 3bpDel (cyan), 15bpDel (magenta), and 39bpDel (orange) rice MAPK4. The green arrow indicates the first residue of the consensus structure among the four models. (B) 3D-alignment of the N termini to the 50^th^ residue of the wild-type sequence. Yellow arrows, β-sheets; blue lines, turns; red lines, α-helices. Residues surrounded by a red rectangle are the residues shown by the arrow in (A).

**Fig. 4.**
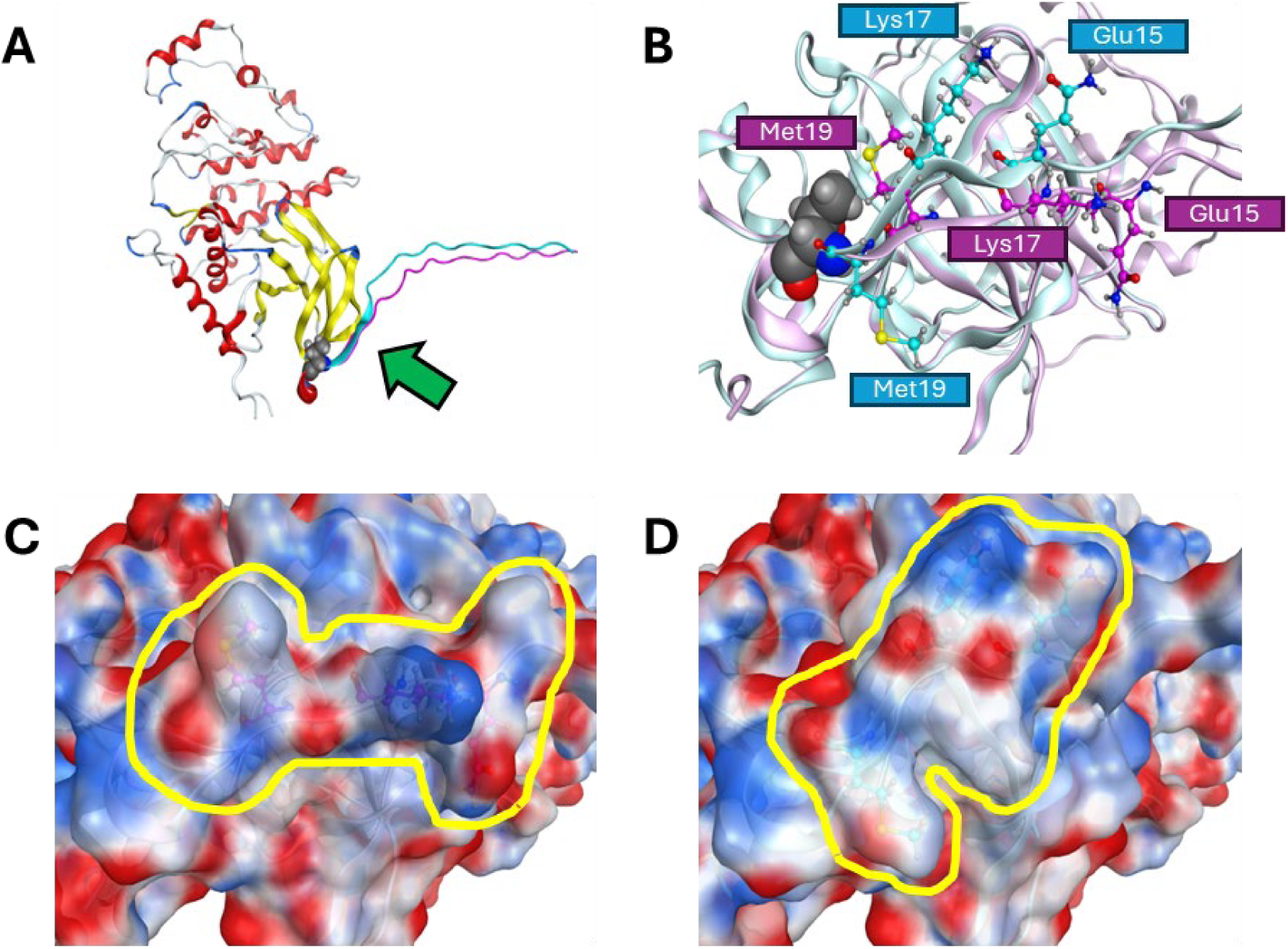
Modeled structures of wild type and Thr21 deletion mutant (3bpDel) of rice MAPK4. (A) 3D-superposition of whole structures. The identical parts of the sequences are shown as ribbons using the same color code as in Fig. 3A, and the differences are indicated by magenta and cyan ribbons for the wild type and the mutant, respectively. (B) Structure viewed from the direction of the green arrow in (A). The residue represented by a space-filling model is Thr21 in the wild-type sequence. The pink ribbon and magenta ball-and-stick models indicate wild type; the light blue ribbon and cyan models indicate 3bpDel. The balls represent the following atoms: cyan and magenta, carbon; blue, nitrogen; yellow, sulfur; red, oxygen; and gray, hydrogen. (C, D) Molecular surfaces of WT (C) and 3bpDel (D) on the structure of (B). Surfaces are colored according to electrostatic condition (red, blue, and white are negative, positive, and neutral, respectively). Depending on the orientation of the side chains of the Glu15, Lys17, and Met19 residues, the surface shapes (surrounded by the yellow outlines) are greatly different.

Comparison of the genomic sequences of *OsMPK4^Takanari^* and *OsMPK4^Koshihikari^* revealed many polymorphisms in the promoter region, 14 polymorphisms in introns, a 6-bp insertion/deletion (In/Del) in the 5′ UTR, and no polymorphism in the coding region (Fig. 5). DNA marker E was developed within a 239-bp In/Del in the promoter region (Fig. 5). There were eight polymorphisms in the interval between marker E and the start codon, and a candidate for a functional polymorphism was delimited in this interval. According to PLACE, a database of plant *cis*-acting regulatory DNA elements (Higo *et al*., 1999), there are three TATA-binding-protein interaction sites (TATTTAA) (Zhu *et al*., 2002) in the interval from marker E to the start codon in the *OsMPK4^Koshihikari^* promoter. In contrast, there are only two in *OsMPK4^Takanari^* because one of the interacting sites is contained within a 25-bp deletion in Takanari relative to Koshihikari (Fig. 5).

**Fig. 5.**
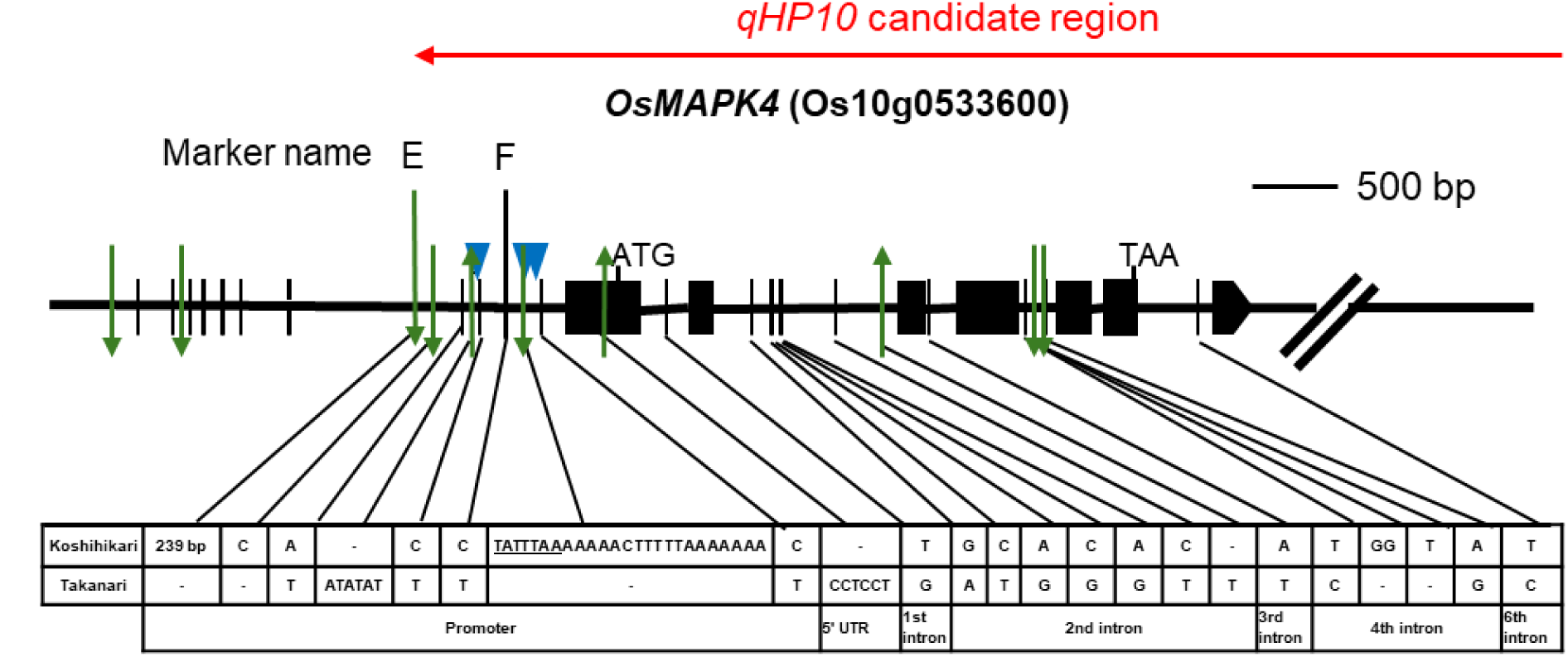
Comparison of genomic sequences of *OsMPK4^Koshihikari^* and *OsMPK4^Takanari^*. Exons and introns are shown as black boxes and horizontal lines, respectively. Vertical lines represent single-nucleotide polymorphisms (SNPs). Green arrows indicate insertions (up) and deletions (down) in Koshihikari relative to Takanari. Blue triangles show the TATA-binding-protein interaction sites (TATTTAA) in Koshihikari; Takanari is missing the second of the three sites owing to the indicated 25-bp deletion.

Figure 6 shows the distribution of the 25-bp In/Del among cultivated and wild rice according to the TASUKE+ browser for the NARO Genebank Core Collection (Kumagai *et al.,* 2019; Tanaka *et al*., 2020; http://ricegenome-corecollection.dna.affrc.go.jp/). Based on ecology and genetics, *O. sativa* is broadly classified into *indica*, *aus*, temperate *japonica*, tropical *japonica*, and aromatic groups (Glaszmann, 1987; Garris *et al*., 2005; Zhao *et al*., 2011; Wang *et al*., 2018). The Takanari-type 25-bp deletion was found in 93.8%, 50%, 6.4%, 8.1%, and 100% of accessions from the *indica*, *aus*, temperate *japonica*, tropical *japonica*, and aromatic group, respectively. Among the accessions of wild species, the deletion occurred in 4.7% of *O. rufipogon* accessions and 80% of *O. nivara* accessions.

**Fig. 6.**
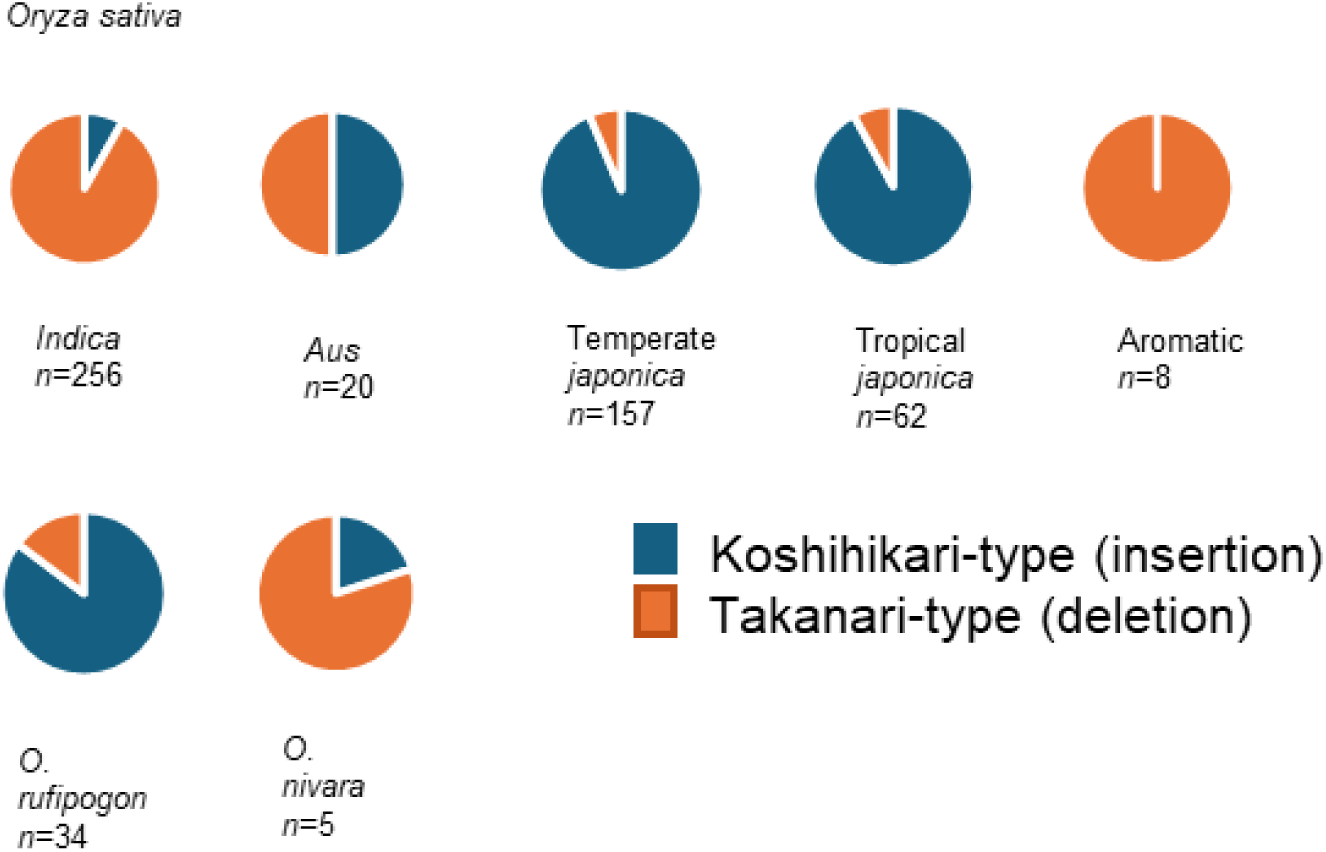
Distribution of the 25-bp In/Del among cultivated and wild rice accessions.

### Characterization of NIL-*OsMPK4*

Based on the previous results, we developed NIL-*OsMPK4,* which carries a <80.2-kb segment containing the *OsMPK4^Takanari^*allele in the Koshihikari genetic background (Fig. 7A). Koshihikari and NIL-*OsMPK4* had nearly identical heading dates, differing by less than one day (Fig. 7B). At the full heading stage, the mRNA level of *OsMPK4* was lower in NIL-*OsMPK4* than in Koshihikari (Fig. 7C). Conversely, leaf photosynthetic rate and stomatal conductance were higher in NIL-*OsMPK4* than in Koshihikari (Fig. 7D, E), whereas the internal CO_2_ concentration and SPAD values were similar between the two lines (Fig. 7F, Supplementary Fig. S3A).

**Fig. 7.**
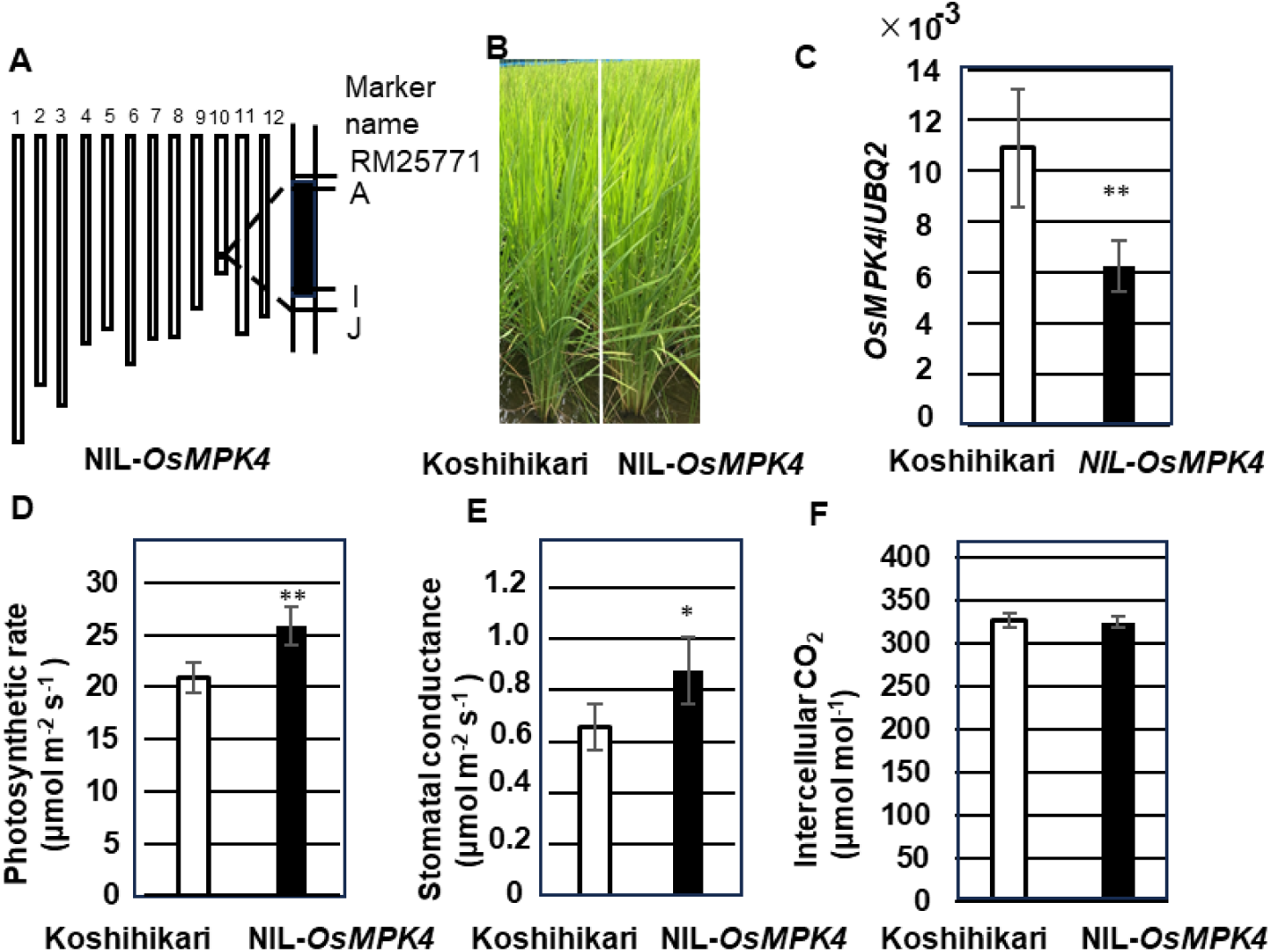
Phenotypic comparison between Koshihikari and NIL-*OsMPK4*. (A) Graphical genotype of NIL-*OsMPK4*. Black rectangle, chromosome region homozygous for Takanari allele; white boxes, homozygous Koshihikari background. (B) Plants at heading stage. (C) Relative expression level of *OsMPK4* in flag leaves at full heading stage. Error bars indicate SD (*n*=5). (D) Photosynthetic rate. (E) Stomatal conductance. (F) Internal CO_2_ concentration. Error bars in (D–F) indicate SD (*n*=6). ** *P*<0.01 and * *P*<0.05 compared with Koshihikari, Student’s *t*-test.

The difference in photosynthetic rate between Koshihikari and NIL-*OsMPK4* increased with irradiation increase, whereas the response to CO_2_ was similar between the two lines (Fig. 8A, B). The maximal rate of ribulose-1,5-bisphosphate (RuBP) carboxylation by Rubisco (*V*_cmax_) and the maximal electron transport rate (*J*_max_), both estimated from the CO_2_ response curve, were also comparable between the lines (Supplementary Fig. S3B, S3C). Anatomical analysis revealed that stomatal length and stomatal densities in the abaxial epidermis were similar between the lines (Supplementary Fig. S4A, S4B).

**Fig. 8.**
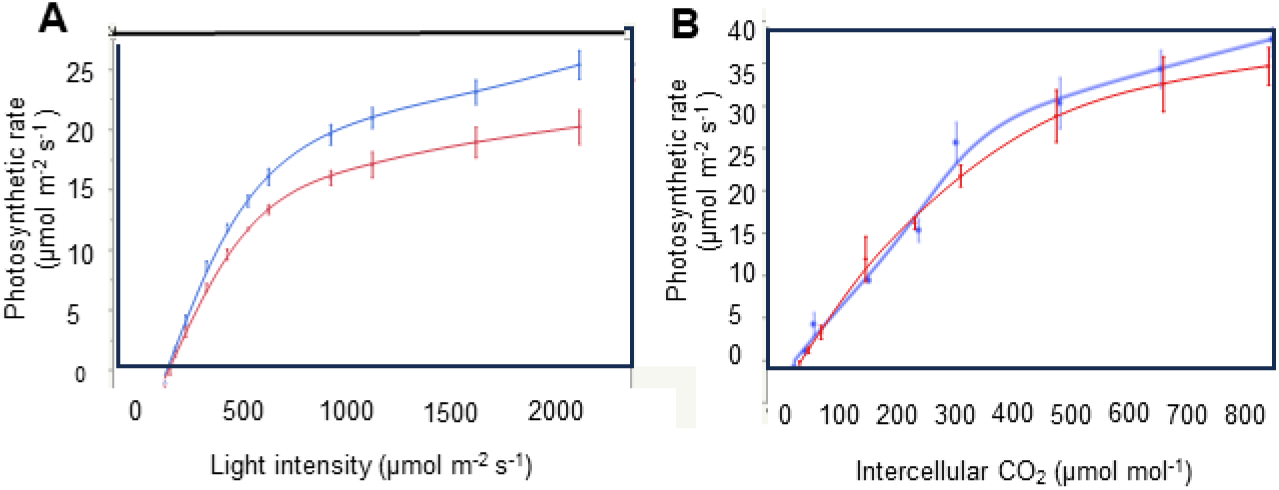
Gas exchange measurements of Koshihikari (red) and NIL-*OsMPK4* (blue). (A) Light response curves. (B) CO_2_ response curves. Data used in the graphs represents averages for Koshihikari and NIL-*OsMPK4*. Error bars indicate SD (*n*=3)

### Effect of *OsMPK4* on agronomic traits

Table 1 compares agronomic traits between Koshihikari and NIL-*OsMPK4* grown in NARO Kannondai Field. Plant height (PH) and panicle number per plant (PNP) were significantly higher in NIL-*OsMPK4* than in Koshihikari, whereas yield parameters (grain number per panicle, fertility ratio, paddy rice weight, ungraded brown rice weight, and graded brown rice weight) and grain quality traits (perfect grain ratio and 1000-grain weight) did not differ significantly between the lines (Table 1). Although the measured biomass production (total weight) of NIL*-OsMPK4* was 11% higher than that of Koshihikari, this difference did not reach statistical significance; nevertheless, this trait may warrant further investigation.

**Table 1.**
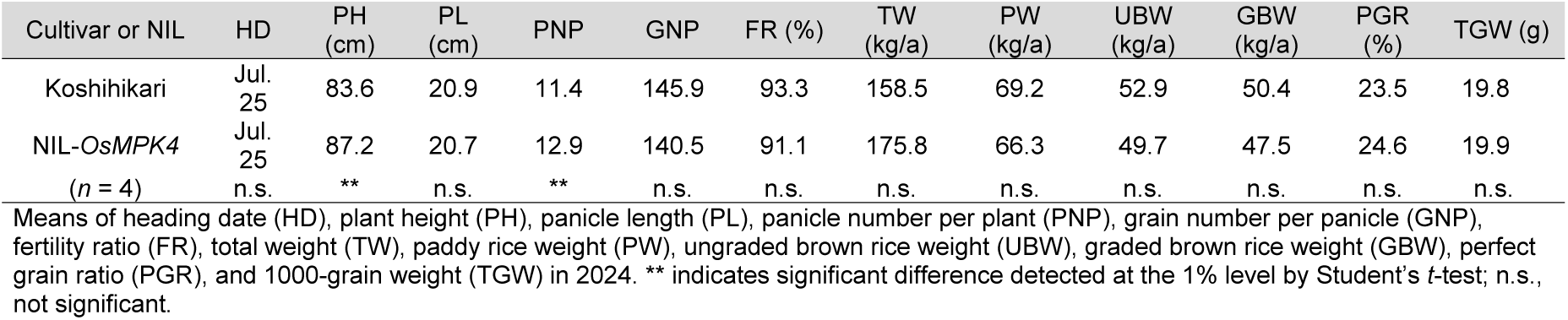
Agronomic traits of Koshihikari and NIL-OsMPK4 grown in the NARO Kannondai field.

| Cultivar or NIL | HD | PH (cm) | PL (cm) | PNP | GNP | FR (%) | TW (kg/a) | PW (kg/a) | UBW (kg/a) | GBW (kg/a) | PGR (%) | TGW (g) |
| --- | --- | --- | --- | --- | --- | --- | --- | --- | --- | --- | --- | --- |
| Koshihikari | Jul. 25 | 83.6 | 20.9 | 11.4 | 145.9 | 93.3 | 158.5 | 69.2 | 52.9 | 50.4 | 23.5 | 19.8 |
| NIL-*OsMPK4* | Jul. 25 | 87.2 | 20.7 | 12.9 | 140.5 | 91.1 | 175.8 | 66.3 | 49.7 | 47.5 | 24.6 | 19.9 |
| (n = 4) | n.s. | ** | n.s. | ** | n.s. | n.s. | n.s. | n.s. | n.s. | n.s. | n.s. | n.s. |

Means of heading date (HD), plant height (PH), panicle length (PL), panicle number per plant (PNP), grain number per panicle (GNP), fertility ratio (FR), total weight (TW), paddy rice weight (PW), ungraded brown rice weight (UBW), graded brown rice weight (GBW), perfect grain ratio (PGR), and 1000-grain weight (TGW) in 2024. \*\* indicates significant difference detected at the 1% level by Student's *t*-test; n.s., not significant.

It has been reported that suppression of *OsMPK4* transcripts enhances rice resistance to bacterial blight (Yuan *et al*., 2007). To test this in our materials, we inoculated Koshihikari and NIL-*OsMPK4* with Japanese *Xanthomonas oryzae* pv. *oryzae.* No significant difference in lesion length was observed between the two lines (Supplementary Fig. S5).

## Discussion

In a study under field conditions, we identified *OsMPK4* as the causal gene underlying the high-photosynthesis QTL *qHP10* and revealed its role in regulation of photosynthesis. Using map-based cloning and CRISPR/Cas9-mediated validation, we demonstrated that reduced *OsMPK4* expression enhances stomatal aperture and CO_2_ uptake, thereby enhancing photosynthesis. A NIL carrying the *OsMPK4^Takanari^* allele (NIL-*OsMPK4*) exhibited a 15–25% increase in photosynthetic rate and stomatal conductance compared with Koshihikari. These findings reveal a mechanism linking MAP kinase signaling to stomatal regulation and carbon assimilation in rice, suggesting new genetic strategies for improving crop productivity. Below, we discuss the physiological implications of this mechanism and its potential application in breeding programs.

### Map-based cloning of *qHP10*

By high-resolution mapping, we delimited the *qHP10* region on chromosome 10 to a 14.5-kb genomic region carrying the MAP kinase gene *OsMPK4* (Fig. 1B). Genome-editing mutagenesis confirmed that *OsMPK4* s the causative gene for this QTL, as reduced expression increased photosynthetic rate under greenhouse conditions (Fig. 2B). In NIL-*OsMPK4*, which was tested under field conditions, the increase in photosynthetic rate was associated with higher stomatal conductance (Fig. 7D, E). These results indicate that MAP kinase signaling influences gas exchange and carbon assimilation under field conditions—a mechanism that could be exploited to improve photosynthetic efficiency in breeding programs.

MAPK cascades are conserved signaling pathways in transducing extracellular stimuli into cellular responses in eukaryotes (Nakagami *et al*., 2005). They consist of an MAP kinase kinase kinase (MAPKKK), an MAP kinase kinase (MAPKK), and an MAP kinase (MAPK) (Mishra *et al*. 2006). MAPKs phosphorylate a variety of substrates such as transcription factors, protein kinases, cytoskeleton-associated proteins, and transporters (Rodriguez *et al*., 2010). Several reports showed the relation between MAPKs and photosynthesis. In *Arabidopsis*, MPK3 and MPK6 activation rapidly alters the expression of photosynthesis-related genes and inhibits photosynthesis during effector-triggered immunity (Su *et al*., 2018). In a recent study, rice lines overexpressing *OsMPK3* showed lower net assimilation rate and reduced stomatal conductance relative to wild-type controls, whereas lines overexpressing *OsMPK6* had higher net carbon assimilation rate and stomatal conductance; in each case these changes resulted from regulation of nuclear-encoded photosynthetic genes (Jonwal *et al*., 2023). In potato (*Solanum tuberosum* L.), *StMPK11* overexpression promoted photosynthesis under drought conditions (Zhu *et al*., 2021). Because *MPK4* is involved in the regulation of stomatal aperture in *Arabidopsis* and tobacco (Hõrak *et al*., 2016; Lin and Chen, 2018), we considered *OsMPK4* as the most likely candidate gene in the 14.5-kb region delimited by studies of recombinants in the *qHP10* region.

The comparison of nucleotide sequence between *OsMPK4^Koshihikari^*and *OsMPK4^Takanari^* revealed three TATA-binding-protein interaction sites (TATTTAA) (Zhu *et al*., 2002) in a delimited region of the *OsMPK4^Koshihikari^* promoter but only two in the same region of the *OsMPK4^Takanari^* promoter (Fig. 5). Furthermore, *OsMPK4* is homologous to tobacco *MPK4* (*NtMPK4*) (Yanagawa *et al*., 2016), and *NtMPK4*-silenced plants have higher photosynthetic rate and stomatal conductance than wild-type plants (Gomi *et al*., 2005). Considering the reduced number of TATA-binding-protein interaction sites in the promoter of *OsMPK4^Takanari^*, we propose that *OsMPK4* expression is lower in Takanari than in Koshihikari, although this remains to be confirmed experimentally.

As shown in Fig. 6, *indica* and *japonica* subspecies had both Takanari- and Koshihikari-type alleles of the 25-bp In/Del containing the TATA-binding-protein interaction site. This suggests that ancestral populations of cultivated rice harbored both alleles. Moreover, most *O. rufipogon* accessions had the Koshihikari-type (insertion) allele, whereas most of the *O. nivara* accessions carried the Takanari-type (deletion) allele. A similar pattern was observed in *japonica* vs. *indica* accessions, respectively, consistent with the fact that *O. rufipogon* is regarded as the progenitor of *japonica*, whereas *O. nivara* is regarded to be the progenitor of *indica* (Garris *et al*., 2005).

In the genome-editing study, the inability to obtain homozygous *OsMPK4* knockout mutants was consistent with the findings of Minkemberg *et al*. (2017), which reported that MAP kinase genes are indispensable for normal plant development, suggesting that OsMPK4 plays a critical role in this process. We selected plants heterozygous for wild-type and knockout *OsMPK4* alleles and found that the photosynthetic rate and stomatal conductance of the heterozygous plants was higher than those of control plants (Fig. 2B). We speculate that the photosynthetic rate of the heterozygous mutant plants was increased because *OsMPK4* mRNA expression was reduced by the loss of a functional allele (although this remains to be confirmed experimentally). Taken together, our results suggest that *OsMPK4* controls photosynthetic traits in a semi-dominant manner: crosses between parents containing different active alleles showed photosynthetic rate and stomatal conductance intermediate between those of the parents (Supplementary Fig. S1).

### Effects of *OsMPK4* assessed by using NIL-*OsMPK4*

The photosynthesis rate and stomatal conductance of NIL-*OsMPK4* (carrying *OsMPK4^Takanari^* in a Koshihikari background) were higher than those of Koshihikari (Fig. 7D, E). In contrast, SPAD, *V*_cmax_, and *J*_max_ (Supplementary Fig. S3) were comparable between the two lines. These results suggest that *OsMPK4* primarily functions by contributing to increased stomatal conductance and thereby enhancing CO_2_ uptake, rather than by improving the enzymatic capacity of the photosynthetic machinery. Stomatal conductance is affected by stomatal length, stomatal density, and aperture (Maruyama and Tajima, 1990; Ohsumi *et al*., 2007). Because stomatal length and density did not differ between Koshihikari and NIL-*OsMPK4* (Supplementary Fig. S4), we infer that the observed difference in stomatal conductance is likely attributable to variation in stomatal aperture.

Accumulating evidence across various plant species suggests that reduced MPK4 activity promotes stomatal opening. In tobacco, silencing of *NtMPK4* impaired CO_2_-induced stomatal closure (Marten *et al*. 2008). In *Arabidopsis*, HIGH LEAF TEMPERATURE1 (HT1) kinase negatively regulates high-CO_2_-induced stomatal closure (Hashimoto *et al*., 2006; Hashimoto-Sugimoto *et al*., 2016; Hõrak *et al*., 2016). MPK4 and MPK12 inhibit HT1 kinase activity in guard cells, thereby modulating stomatal responses to CO_2_ (Hõrak *et al*., 2016; Jakobson *et al*., 2016; Tõldsepp *et al*., 2018). In this study, we observed that NIL-*OsMPK4* had higher stomatal conductance and photosynthetic rate than Koshihikari under ambient CO₂ conditions (Fig. 7D, E). Although we measured CO₂ response curves for photosynthesis (Fig. 8B), we did not directly evaluate stomatal CO₂ sensitivity (i.e., the extent of stomatal closure in response to elevated CO₂). Further experiments, such as direct measurements of stomatal aperture or conductance in response to stepwise changes in CO₂ concentration, will be necessary to confirm altered CO₂ sensitivity in NIL-*OsMPK4.* Taken together with our NIL analysis, these findings suggest that partial loss-of-function alleles of *OsMPK4* can enhance photosynthetic performance without causing lethality, likely by modifying stomatal regulation rather than completely abolishing MPK4-mediated signaling.

In genome-editing experiments, plants homozygous for 3-, 15-, and 39-bp deletions of *OsMPK4* could be selected from T_1_ progenies (Fig. 2A). All of these are in-frame mutations and therefore do not disrupt the *OsMPK4* open reading frame. These plants showed higher photosynthesis rate than those containing the wild-type allele (Fig. 2B). Because complete loss of OsMPK4 function is lethal (Minkemberg *et al*., 2017), we assume that these mutated OsMPK4 proteins retain partial functionality.

Recently, Takahashi *et al*. (2022) reported that in *Arabidopsis* under low-CO_2_ conditions, HT1 kinase activates another protein kinase, CONVERGENCE OF BLUE LIGHT AND CO_2_ (CBC1/CBC2), through phosphorylation, thereby inducing stomatal opening. Conversely, under high-CO_2_ conditions, HT1 kinase interacts with MPK4/MPK12, and this interaction inhibits HT1 kinase activity, resulting in stomatal closure. If a similar mechanism operates in rice, plants homozygous for the 3-, 15-, and 39-bp deletions in *OsMPK4* might produce mutated OsMPK4 proteins with reduced ability to interact with rice HT1 kinase. This could lead to a partial loss of CO_2_-induced stomatal responses and impaired stomatal closure under ambient CO_2_ conditions. Recently, *OsHT1* (Os06g0636600) was identified and its product was found to interact with *Arabidopsis* MPK12, causing stomatal closure (Xiao *et al*., 2024). Therefore, it is necessary to determine whether rice HT1 kinase interacts with wild-type and/or mutated OsMPK4.

### Agronomic trait evaluation

In the field comparison between Koshihikari and NIL-*OsMPK4*, we found no significant differences in yield or grain quality (Table 1). The increased plant height and panicle number per plant of NIL-*OsMPK4* (Table 1) might contribute to its biomass, although the increase measured in our study was not statistically significant. In our previous study, NIL10 produced 6–10% more biomass at harvest and 8–14% higher grain yield than Koshihikari (Yamashita *et al*., 2022). NIL10 carries a 2.8 Mb segment from Takanari, whereas NIL-*OsMPK4* carries a <80.2-kb segment (Fig. 7A). The larger segment in NIL10 may include additional QTLs, such as those related to sink size, that are absent in NIL-*OsMPK4*. By using a NIL with a delimited chromosomal segment, we were able to estimate the specific effect of a single gene. To further increase rice yield, however, it is necessary not only to increase photosynthetic rate (source capacity) but also to expand the sink size and enhance the carbon partitioning to grains (Ueda *et al*., 2025).

Transgenic rice plants in which *OsMPK4* has been silenced by RNAi have enhanced resistance to bacterial blight (Yuan *et al*., 2007). Thus, we hypothesized that NIL-*OsMPK4* would show higher resistance to bacterial blight than Koshihikari. Although transcript levels of *OsMPK4* were reduced in NIL-*OsMPK4* relative to Koshihikari (Fig. 7C), the resistance of NIL-*OsMPK4* was not significantly different from that of Koshihikari (Supplementary Fig. S5). This result suggested that the difference in *OsMPK4* expression between Koshihikari and NIL-*OsMPK4* was insufficient to affect resistance to bacterial blight.

Considering other QTLs for photosynthesis rate such as *GPS* (Takai *et al*., 2013) and *CAR8* (Adachi *et al*., 2017), NIL-*GPS*, which carries a segment including the Takanari allele of *GPS* in the Koshihikari genetic background, had a clearly increased photosynthetic rate per unit leaf area, but its narrow leaves (small leaf area) reduced the photosynthesis capacity on the whole-leaf level (Hirotsu *et al*., 2017). The yield of NIL-*GPS* was similar to that of Koshihikari (Takai *et al*., 2013; Ueda *et al*., 2021). *CAR8* encodes the Heme Activator Protein 3 (HAP3) subunit of a CAAT-box-binding transcription factor called OsHAP3H (Wei *et al*., 2010; Yan *et al*., 2011). Adachi *et al*. (2017) studied the characteristics of NIL(*CAR8*), which carries a segment including the Habataki *CAR8* allele in the Koshihikari genetic background, because Habataki is a high-yielding *indica* cultivar widely used in QTL studies for traits related to sink size and photosynthetic capacity. This NIL has increased flag leaf nitrogen content, stomatal conductance, and photosynthesis but also an accelerated heading date, resulting in low yield (Adachi *et al*., 2017). On the other hand, *qHP10*/*OsMPK4^Takanari^* enhances stomatal conductance, resulting in high photosynthesis rate, without any detected negative pleiotropic effects. This gene, one of the few source capacity genes that have been identified, increases source capacity by 15–25%, and we expect to increase yield by pyramiding with a gene related to sink size.

## Supplementary data

**Supplementary Table S1.** Primers used in this study.

**Supplementary Fig. S1.** Identification of *qHP10*.

**Supplementary Fig. S2.** Fine mapping of *qHP10*.

**Supplementary Fig. S3.** Photosynthesis-related traits in Koshihikari and NIL-*OsMPK4*.

**Supplementary Fig. S4.** Observation of stomata in Koshihikari and NIL-*OsMPK4*.

**Supplementary Fig. S5.** Length of the lesions resulting from bacterial blight infection of Koshihikari and NIL-*OsMPK4*.

## Acknowledgements

We acknowledge Dr. J. Wu and Dr. Y. Katayose for help with the Takanari genome sequence analysis. We are grateful to Prof. T. D. Sharkey for kindly sending us an A–Ci curve-fitting utility based on Microsoft Excel program ver. 2.95. We are also grateful to Dr. T. Tanabata for kindly sending us his original computer software to count stomatal numbers. We thank Mrs. M. Suzuki and Mrs. M. Iizumi for their technical assistance. We also thank Dr. K. Hori for instructions on qRT-PCR. Computer resources were supported by Agriculture, Forestry and Fisheries Research Information Technolofgy Center, AFFRC.

## Author contributions

TU, TY: conceptualization; TU, SA, TH, and TY: methodology; MHM: formal analysis; TU, SA, UY, RM, and JT: investigation; KS: resources (CRISPR/Cas9-mediated mutant plants); TU, SA, MHM, UY, RM, YT, TH, TY, and JT: writing – original draft; All authors: writing – review & editing; YT and JT: supervision; TU, SA, and TY: funding acquisition.

## Data Availability

Sequence data from this study have been deposited with the DDBJ/EMBL/GenBank databases under accession no. LC867645.

