## Supplementary Figure / Table for "Natural variation in rice *mitogen-activated protein kinase 4* contributes to increased photosynthetic rate under field conditions"

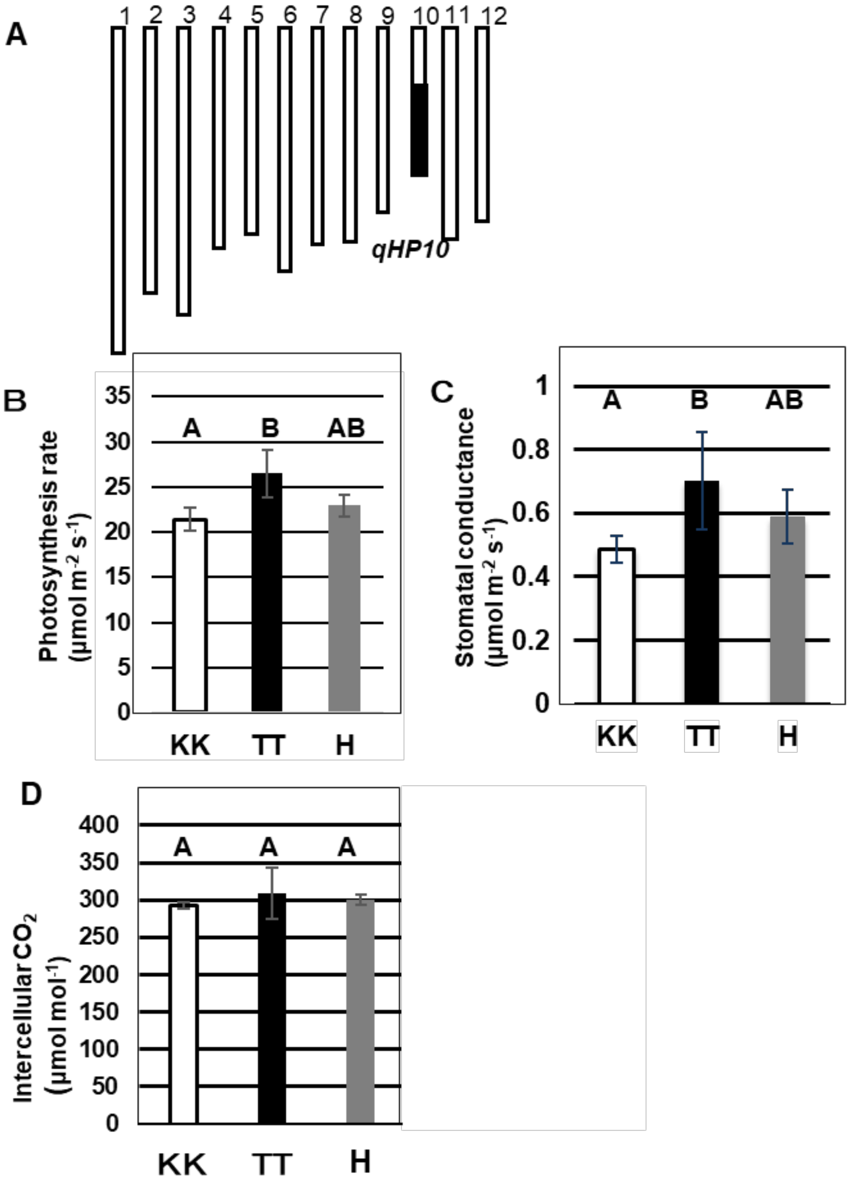


**Supplementary Fig. S1.** Identification of *qHP10*. (A) Graphical genotype of SL1235 (Adachi *et al*., 2019). Black box, homozygous chromosome region from Takanari; white boxes, homozygous Koshihikari background. (B–D) Comparison of the photosynthetic rate (B), stomatal conductance (C), and internal $CO_2$ concentration (D). KK, homozygous for Koshihikari *qHP10* allele; TT, homozygous for Takanari *qHP10* allele; H, heterozygous. Error bars indicate SD ($n$=8). Values followed by the same letters indicate no significant difference among rice lines at $P$<0.05 by LSD test.

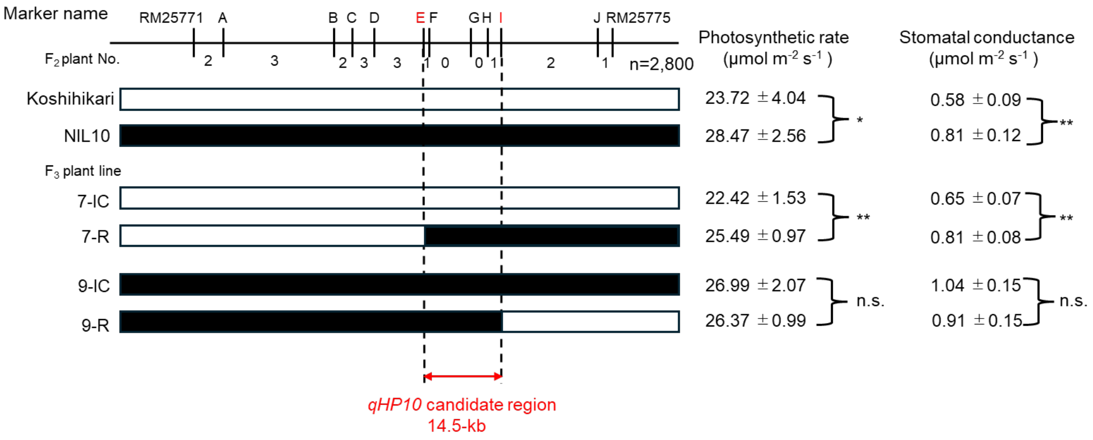


**Supplementary Fig. S2.** Fine mapping of *qHP10.* Ten DNA markers were used for genotyping. White and black rectangles indicate homozygous regions for Koshihikari and Takanari chromosome segments, respectively. Koshihikari and NIL10 were used as controls. On the right, the photosynthetic rate and stomatal conductance of flag leaves (means ±SD; $n$=6) are shown. ** $P$<0.01 and * $P$<0.05 for comparison between IC (internal control) and R (recombinant) progeny of the same $F_2$ plant using Student's $t$-test. ns., no significant difference

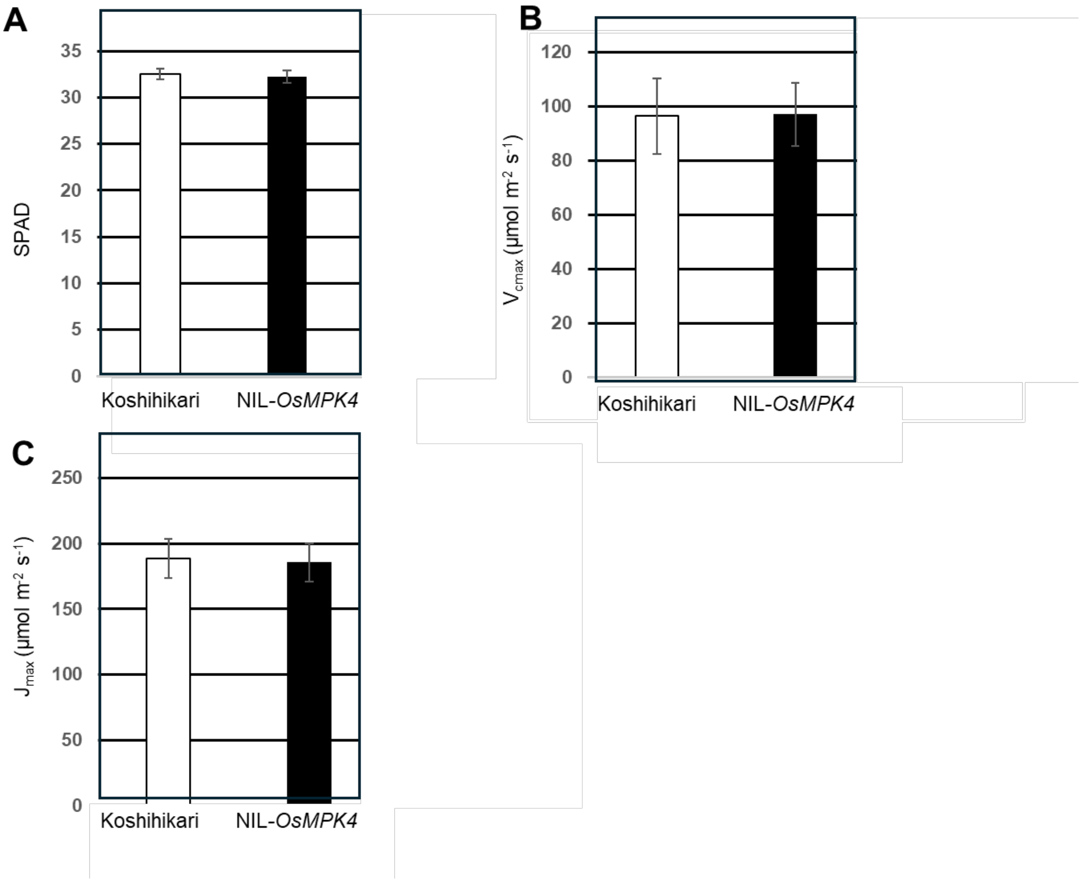


**Supplementary Fig. S3.** Photosynthesis-related traits in Koshihikari and NIL-*OsMPK4.* (A) SPAD. Error bars indicate SD (*n*=6). (B, C) $V_{\text{cmax}}$ (B) and $J_{\text{max}}$ (C) were calculated by fitting an FvCB model with the $CO_2$ response curves measured. Error bars indicate SD (*n*=3). $V_{\text{cmax}}$, maximal rate of ribulose-1,5-bisphosphate (RuBP) carboxylation by Rubisco; $J_{\text{max}}$, maximal photosynthetic electron transport rate

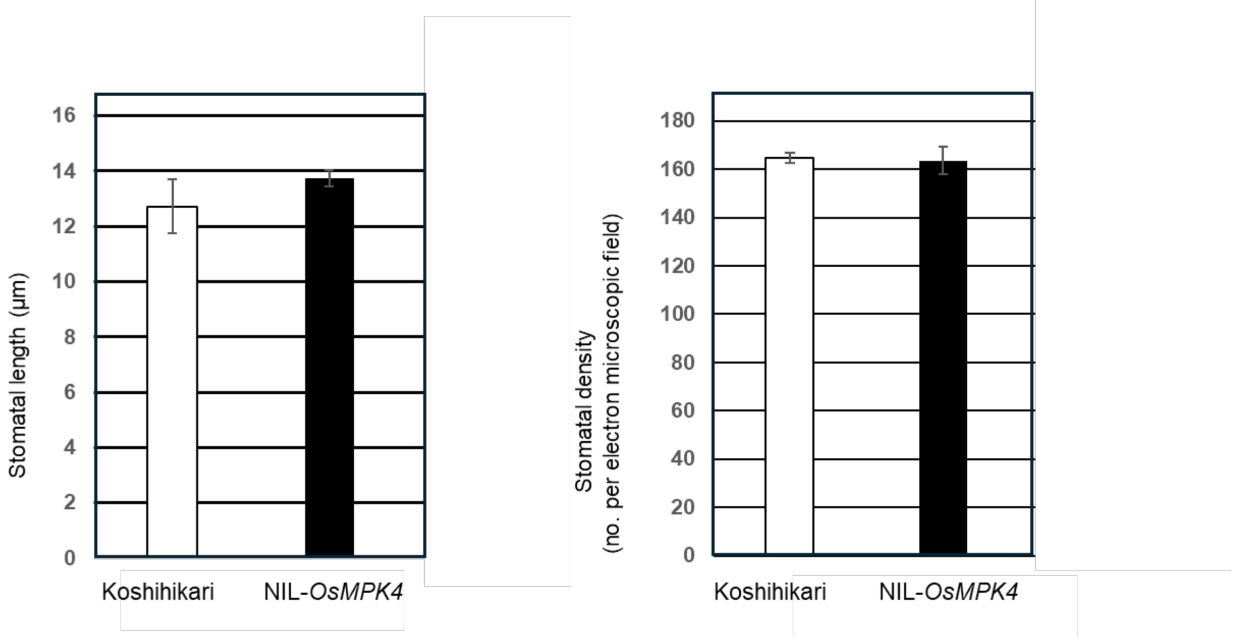


**Supplementary Fig. S4.** Observation of stomata in Koshihikari and NIL-*OsMPK4*. (A) Stomatal length and (B) stomatal density in flag leaves of field-grown plants at full heading. Error bars indicate SD (*n*=3).

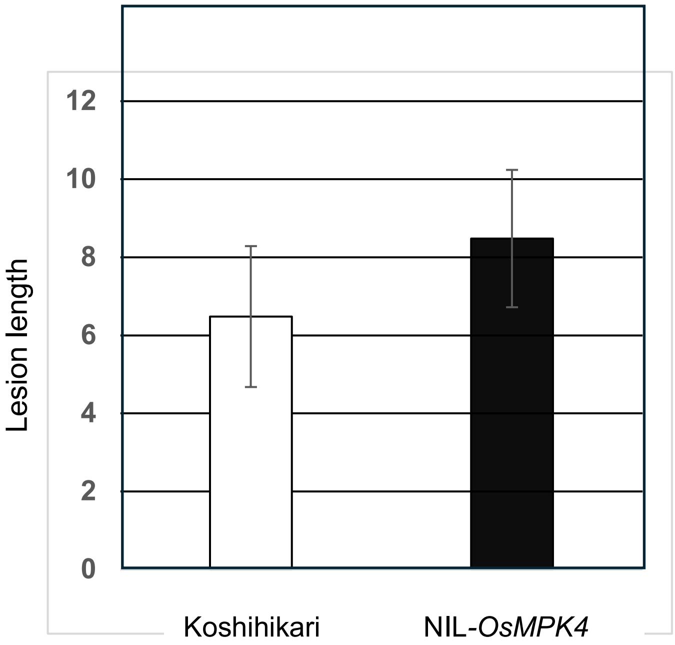


**Supplementary Fig. S5.** Length of the lesions resulting from bacterial blight infection of Koshihikari and NIL-*OsMPK4*. Error bars indicate SD (*n*=6).

Supplementary Table S1. Primers used in this study.

| Marker | Class | U/L | Primer sequence | Restriction enzyme |
|---|---|---|---|---|
| Fine mapping | | | | |
| A | CAPS | U | GAACGCCGCCTAATTAACAA | *Alu* I |
| | | L | ACGAAGGCTTTTGGAGTTTG | |
| B | CAPS | U | CGCGGGGAGCTACTGAAG | *Sma* I |
| | | L | CCTCGAAGTCGATGAACACC | |
| C | In/Del | U | CTTCGGTGGCAAGAGTAGTACC | - |
| | | L | CAACGTGGAATTCTCACCACTA | |
| D | CAPS | U | TTTGAAGAGGGAGGTAGCAG | *Xsp* I |
| | | L | AGCCTCAAAACGGATACGC | |
| E | In/Del | U | CGGCTTGGGACGAAAAGCTC | - |
| | | L | CCTGGTGGAGATTTCGGGC | |
| F | CAPS | U | GCCCGAAATCTCCACCAGG | *Mlu* I |
| | | L | GAATGGGGATCGGAGCGAG | |
| G | In/Del | U | CAAACCTAATACTAGGATGGGACA | - |
| | | L | GAACCAGAAGACAATTGTTTGGA | |
| H | dCAPS | U | ACTTAGCCTAAATTTGATATAATGCTCAATT | *Mfe* I |
| | | L | GCTATCTAATAGTAAAAGTACCAAC | |
| I | CAPS | U | AGTTCATGTCCATAGCGGCT | *Hha* I |
| | | L | AATACAATCGCCCTCCCACG | |
| J | CAPS | U | TCAATGGTGATGCCATTTTG | *Afa* I |
| | | L | TTGAATCGCCATTTCCATTT | |
| Detection of CRISPR/CAS9 induced mutation | | | | |
| Target1 | - | U | AAGCTTCTCCCACCATAGTCCT | - |
| | | L | AGACCTCGAAGAAGTTCCCGTA | |
| Target2 | - | U | GGCCCAAAAATGTTATGGTTT | - |
| | | L | TAGTGGTAAGGCCAAGTTCGAT | |
| qRT-PCR | | | | |
| OsMPK4 | - | U | TTGTGACTCGTCAACCCCTG | - |
| | | L | TGAGTCATCTGGCGACCCTA | |
| UBQ2 | - | U | GAGCCTCTGTTCGTCAAGTA | - |
| | | L | ACTCGATGGTCCATTAAACC | |
